# ExoShorkie: Predicting RNA-seq coverage of exogenous genomes in yeast by transfer learning

**DOI:** 10.64898/2026.01.25.701486

**Authors:** Jonathan Mandl, Yaron Orenstein

## Abstract

**Motivation:** Predicting the RNA-seq coverage of native and exogenous sequences is central to many endogenous and synthetic-biology applications. Substantial progress has been made in developing methods to predict the RNA-seq coverage of native genomic sequences, with the recently developed Shorkie achieving state-of-the-art performance in yeast. However, prediction performance of these methods over the challenging out-of-distribution exogenous DNA is still unknown. Recent studies measured RNA-seq coverage of large exogenous genomes in yeast, providing a unique opportunity to train machine-learning models on a large exogenous sequence space and to improve both prediction performance and our understanding of regulatory mechanisms.

**Results:** Here, we introduce ExoShorkie, a method we developed by extending Shorkie through transfer learning across multiple exogenous RNA-seq datasets. We demonstrate that ExoShorkie significantly improves prediction performance on held-out exogenous genomes and outperforms both a native-genome-trained Shorkie baseline and Yorzoi, the only competing method in predicting exogenous RNA-seq coverage in yeast, in cross-validation and in leave-one-genome-out evaluations. Furthermore, through interpretability analyses we reveal biologically meaningful regulatory motifs and distinct regulatory rules in exogenous genomes in yeast, providing new insights into transcriptional regulation beyond native genomic contexts.

**Availability and implementation:** ExoShorkie is available at https://github.com/OrensteinLab/ExoShorkie.

## 1. Introduction

Models that can reliably predict RNA sequencing (RNA-seq) coverage directly from DNA sequence hold promise for enabling the design of synthetic sequences with desired expression levels across a wide range of biotechnological and therapeutic applications. Recent advances in high-throughput genomics and deep learning have driven the development of models that achieve high predictive performance in mapping genomic sequences to gene expression. These models, including Enformer and Borzoi, commonly employ transformer-based neural networks to predict RNA-seq coverage and related genome-wide functional tracks directly from DNA sequence at high resolution [Avsec et al., 2021, Chao et al., 2025, Avsec et al., 2025, Linder et al., 2025].

Despite the high predictive performance these models achieve on native genomic sequences, they have been shown to generalize poorly to out-of-distribution synthetic sequences involving rearrangements of regulatory elements [Ribeiro-dos Santos and Maurano, 2025]. High predictive performance in such a task is essential for the design of complex synthetic sequences with multiple regulatory elements. However, native genomes lack the sequence diversity required for models to learn the full complexity of the gene-regulatory grammar [de Boer and Taipale, 2024]. Consequently, training beyond native genomic sequences may be required to learn complex regulatory rules. Randomized DNA has been proposed as a potential solution to enable models to be trained on a much larger sequence space while reducing the biases and homology inherent to native genomes [de Boer and Taipale, 2024].

While deep-learning models can successfully predict the expression of random promoter sequences in massively parallel reporter assays [Vaishnav et al., 2022, de Boer et al., 2020], these models focus on isolated regulatory elements within short reporter constructs. Consequently, they cannot account for multiple regulatory elements acting jointly within a gene-regulatory architecture [Zrimec et al., 2020]. Computational methods thus remain limited in their ability to predict the genome-scale function of complex synthetic constructs [James et al., 2025]. Training on long random DNA integrated within a chromosomal locus can provide models with examples beyond the native genome of multiple regulatory elements acting together, as well as locus-level mechanisms, such as 3D looping, insulation, and competition [de Boer and Taipale, 2024]. However, generating long random DNA sequences and inserting them into living cells has until recently been considered a highly challenging task [de Boer and Taipale, 2024]. Only recently, experimental techniques have advanced to produce complete synthetic genomes and insert them into living cells [James et al., 2025]. As a result, multiple studies measured the expression of long exogenous DNA sequences in yeast, which can serve as a proxy for the vast space of out-of-distribution DNA [Luthra et al., 2024, Zhou et al., 2022, Camellato et al., 2024, Meneu et al., 2025].

Recent efforts have begun to explore genome-wide RNA-seq coverage prediction in yeast. Shorkie was developed as a transformer-based DNA language model pretrained with a self-supervised objective on diverse fungal genomes and fine-tuned to predict thousands of RNA-seq tracks in yeast [Chao et al., 2025]. However, it is trained solely on native genomic sequences and has not been evaluated on exogenous genomes. In contrast, Yorzoi was developed as a similar transformer architecture to predict RNA-seq coverage on exogenous human DNA sequences in yeast [Schneider et al., 2025]. While the Yorzoi study demonstrated that exogenous DNA can be modeled, it remains unclear whether models trained on one exogenous genome can generalize across other exogenous genomes. Moreover, none of these studies provided interpretability analyses of the regulatory logic captured in exogenous DNA. Addressing these limitations is essential for evaluating both the generalization ability of such models and the regulatory rules they learn from non-native sequences.

Here, we present *ExoShorkie* to accurately predict the RNA-seq coverage of exogenous genomes in yeast. We first demonstrate that the original Shorkie model, when trained exclusively on native genomic sequences, fails to generalize to exogenous DNA. To address this, we developed ExoShorkie, a model trained by fine-tuning Shorkie on RNA-seq datasets of exogenous DNA sequences in yeast. We show that ExoShorkie outperforms both a native-genome-trained baseline and the competing exogenous DNA approach, Yorzoi, in cross-validation and in leave-one-genome-out evaluations, demonstrating generalization across exogenous contexts. Through an ablation analysis, we also show that Shorkie’s pretraining on fungal and native yeast sequences improves the prediction performance on exogenous DNA. Furthermore, our interpretability analyses reveal that ExoShorkie recovers canonical yeast regulatory motifs in exogenous DNA. Notably, many of these motifs are missed or predicted with opposite effects by models trained on native genomic DNA. Overall, our results suggest that regulatory rules learned from exogenous sequences are generalizable and extend beyond those inferred from the native genome. We expect ExoShorkie to be applied to design synthetic sequences with desired expression levels for various applications.

## 2. Methods

### 2.1. RNA-seq datasets

We utilized publicly available datasets from recently published studies that measured strand-resolved RNA-seq coverage of exogenous DNA in *Saccharomyces cerevisiae* (*S. cerevisiae*). To provide a reference for native expression, we also utilized a strand-resolved RNA-seq dataset of the wild-type *S. cerevisiae* genome from Wery et al. [2023]. A summary of all datasets, including their sequence origin and total length, is provided in Table 1.

**Table 1.**
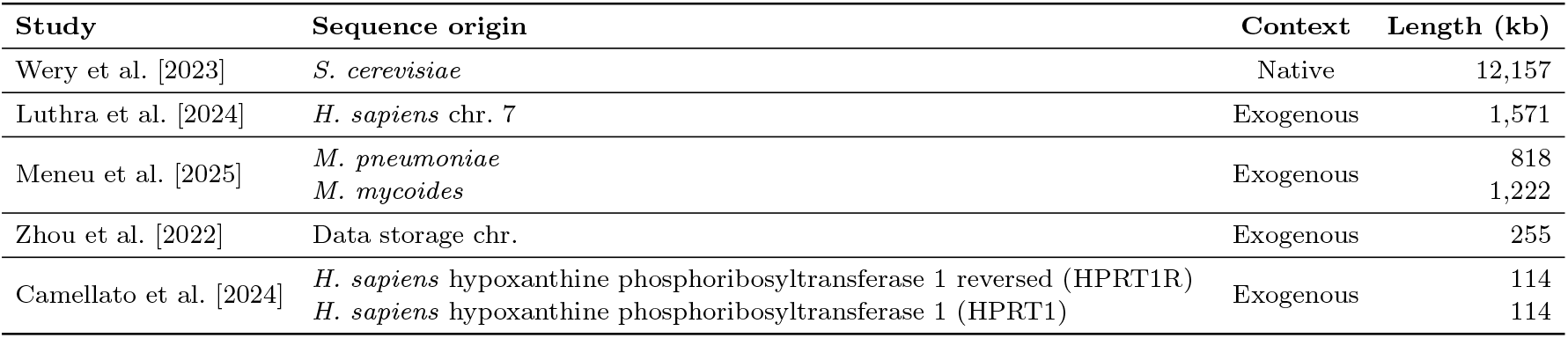
Summary of RNA-seq datasets used for training and evaluation in this study.

| Study | Sequence origin | Context | Length (kb) |
| --- | --- | --- | --- |
| Wery et al. [2023] | *S. cerevisiae* | Native | 12,157 |
| Luthra et al. [2024] | *H. sapiens* chr. 7 | Exogenous | 1,571 |
| Meneu et al. [2025] | *M. pneumoniae* | Exogenous | 818 |
|  | *M. mycoides* |  | 1,222 |
| Zhou et al. [2022] | Data storage chr. | Exogenous | 255 |
| Camellato et al. [2024] | *H. sapiens* hypoxanthine phosphoribosyltransferase 1 reversed (HPRT1R) | Exogenous | 114 |
|  | *H. sapiens* hypoxanthine phosphoribosyltransferase 1 (HPRT1) |  | 114 |

### 2.2. RNA-seq samples pre-processing

Our RNA-seq processing pipeline was adapted from the workflow described by Luthra et al. [2024]. We applied a uniform computational pipeline to all RNA-seq datasets listed in Table 1. We performed initial quality control and adapter removal using Trimmomatic (v0.36) [Bolger et al., 2014]. Specifically, we utilized the `ILLUMINACLIP` module with the TruSeq3-PE-2 adapter library (parameters: 2:30:10:2:true) and a 4 bp sliding window to trim bases where the average Phred quality fell below 10. Reads shorter than 36 bp following trimming were discarded. Post-processing sequence quality was verified using FastQC (v0.11.8) [Simon Andrews et al., 2010].

For the native yeast experiments, we used the yeast S288C reference genome (R64) for read alignment. For the exogenous experiments, we constructed a hybrid reference genome by appending the corresponding exogenous sequence to the yeast S288C (R64) reference. This hybrid reference reflects the mixed genomic content of the cells and prevents spurious alignment of yeast RNA reads to the exogenous sequence.

We conducted sequence alignment using STAR (v2.7.10a) [Dobin et al., 2013]. We indexed the hybrid reference genomes using the parameter `--genomeSAindexNbases` 10. We performed alignments using `--twopassMode Basic, --alignIntronMax 5000`, and `--outFilterType BySJout`. We generated output files as coordinate-sorted BAM files via the `--outSAMtype BAM SortedByCoordinate` parameter. For datasets with multiple replicates, we consolidated BAM files using the Picard tools (v2.18.7) (Broad Institute) `MergeSamFiles` utility and indexed them using Samtools [Li et al., 2009].

To generate the RNA-seq tracks, we utilized the `bamCoverage` module from deepTools (v3.1.3) [Ramírez et al., 2016] to produce strand-specific BigWig files at 1 bp resolution. We normalized the signal using counts per million (CPM) via the `--normalizeUsing CPM`. We generated forward- and reverse-strand tracks separately using the `--filterRNAstrand` flag. To mitigate coverage artifacts arising from high-copy-number repeats, we excluded the rRNA locus on chromosome XII (coordinates chrXII:369,803–499,262) during BigWig generation using the `--blackListFileName` parameter.

### 2.3. ExoShorkie data pre-processing

For each dataset, we segmented the base-resolution RNA-seq coverage tracks into overlapping windows according to the input length of Shorkie: 16,384 bp. We one-hot encoded the resulting windows to fit the expected input of Shorkie. We concatenated this encoding with the species one-hot vector used by Shorkie, a 165-dimensional representation in which the *S. cerevisiae* channel (index 114) is set to 1. To generate the target labels, we followed the pre-processing of the Shorkie study: we cropped 1,024 bp from both ends of the window to mitigate edge effects, retaining the central 14,336 bp. The ground-truth RNA-seq signal was then summed across non-overlapping 16 bp bins, yielding a final target vector of 896 elements per window, consistent with the model’s output architecture.

To ensure the same training set size across all exogenous datasets, despite their varying total lengths, we computed a dataset-specific stride based on the training sequence length *L* and window size *W* , such that slightly more than 10,000 training windows were generated per dataset at this step, with twice that number when including the reverse-complement windows:

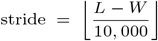

To normalize the target labels during training, we applied a log transformation, log(*x* + 1), to all 896-dimensional target vectors. Subsequently, we performed z-score normalization using a global mean and standard deviation computed across all bins and all windows in the training set. For evaluation, we used the target labels at their original coverage scale, as performance was measured using Spearman correlation, which is invariant to monotonic transformations and depends only on rank ordering.

### 2.4. ExoShorkie model architecture

The Shorkie architecture is a U-Net–transformer–based model that operates on 16,384 bp input sequences [Chao et al., 2025]. Briefly, the input sequence is first processed by a convolutional encoder with progressive downsampling, followed by a stack of transformer layers that capture long-range dependencies. A decoder with U-Net–style skip connections then restores sequence resolution and produces predictions at 16 bp resolution.

We adapted this architecture to create ExoShorkie by retaining the trained Shorkie feature-extraction trunk as a backbone and adding a new task-specific dense output layer to predict the experimental tracks used in this study, while ignoring the outputs of the original Shorkie prediction heads (Fig. 1A).

**Fig. 1.**
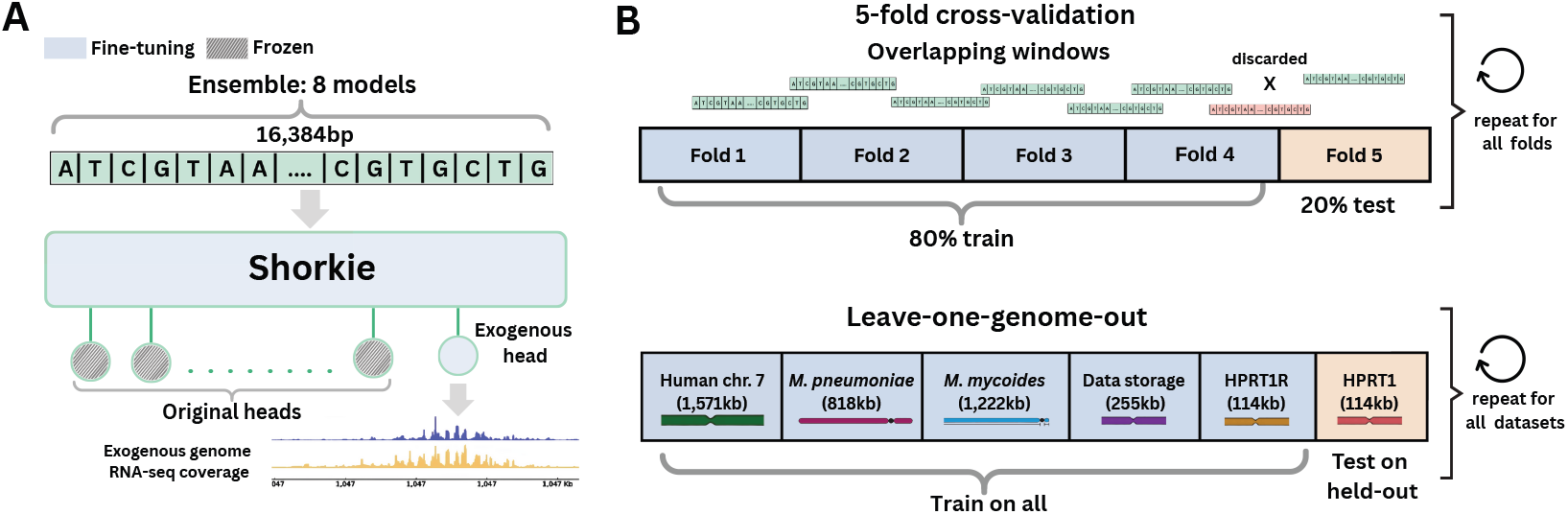
Overview of the Shorkie model fine-tuning and performance evaluation (A) Model fine-tuning scheme. An additional prediction head is added to Shorkie, and both the trunk and the exogenous head are fine-tuned jointly to predict RNA-seq coverage from a 16,384 bp-long exogenous DNA sequence input expressed in yeast cells, while the original Shorkie prediction heads are frozen. To improve robustness, an ensemble of eight models is trained for each cross-validation split. (B) Evaluation strategies. In 5-fold cross-validation, each exogenous genome is partitioned by genomic coordinates into five folds, with one fold used for testing and four for training. Sequences are segmented into overlapping windows, and windows overlapping both training and test folds are discarded. In addition, a leave-one-genome-out evaluation is performed, in which an entire exogenous genome is held out to assess generalization to unseen exogenous DNA.

### 2.5. ExoShorkie training strategy and hyper-parameters

To fine-tune Shorkie, we treated each exogenous dataset as a separate learning task and fine-tuned the model independently on each dataset. We utilized the trained Shorkie feature-extraction trunk, attaching a new task-specific output layer. We then fine-tuned the Shorkie trunk and the new output layer jointly, while ignoring the output of the original task-specific heads (Fig. 1A).

We performed fine-tuning in two stages. First, since the original Shorkie model outputs non–strand-resolved RNA-seq coverage predictions whereas our target data are strand-resolved, we performed a native genomic adaptation phase on native yeast RNA-seq data. We fine-tuned the model on the native Wery et al. dataset for 5 epochs with a batch size of 32 and a learning rate of 2 *·*10^−5^, matching the learning rate used in the original Shorkie fine-tuning. Subsequently, we fine-tuned this adapted model on the specific exogenous dataset for 3 epochs to limit drift from the trained representation, with a batch size of 32 and a learning rate of 1 *·*10^−5^, following learning-rate settings used for Enformer fine-tuning [Ribeiro-dos Santos and Maurano, 2025]. In both stages, we utilized the Adam optimizer (default parameters: *β*_1_ = 0.9, *β*_2_ = 0.999) and minimized the mean-squared-error loss.

All model fine-tuning was performed on a Linux server (Intel Xeon Gold 6338 @ 2.00 GHz, 1*×* NVIDIA A100 80 GB GPU, 512 GB RAM). The full fine-tuning procedure, including all models trained in this study, completed in under 56 hours.

### 2.6. ExoShorkie ensemble learning strategy

We employed an ensemble learning strategy in ExoShorkie to improve prediction robustness, i.e., minimize its variance. The original Shorkie model is distributed as an ensemble of eight distinct model instances, each originally trained on a different split of an 8-fold cross-validation scheme. During the genomic adaptation phase, we fine-tuned each of the eight Shorkie models independently, utilizing distinct random seeds for data shuffling and for the initialization of the task-specific output layers. These adapted models served as the starting point for all subsequent exogenous fine-tuning. During the 5-fold cross-validation on each exogenous dataset, all eight instances were fine-tuned on the training set of each fold, resulting in a total of 40 trained ensemble models per dataset.

For cross-validation evaluation, predictions for each test sequence were obtained by averaging the outputs of the eight ensemble members trained on the corresponding fold. For leave-one-genome-out evaluation, predictions for each held-out genome were computed by averaging the outputs of all ensemble models trained on the remaining exogenous datasets (40 models per dataset).

### 2.7. Prediction performance evaluation

#### 2.7.1. Cross-validation evaluation

We evaluated prediction performance within each dataset using a 5-fold cross-validation strategy (Fig. 1B). To prevent data leakage, we partitioned each exogenous genome into five contiguous non-overlapping genomic intervals based on positional coordinates. In each iteration, four intervals (80%) were utilized for training and one interval (20%) was held out for testing. We discarded any windows spanning the train-test boundary to strictly avoid train-test overlap. To maximize robustness, we trained an ensemble of 8 independently initialized models for each cross-validation fold. Test intervals were segmented into overlapping 16,384 bp windows with a 1,024 bp stride to ensure complete coverage. We quantified prediction performance using the median Spearman correlation coefficient between the predicted and observed coverage tracks over all test windows.

#### 2.7.2. Leave-one-genome-out evaluation

To assess the model’s ability to generalize to unseen exogenous genomes, we employed a leave-one-genome-out strategy (Fig. 1B). For this evaluation, each exogenous dataset served as a held-out test set, while predictions were generated by an ensemble comprising all models trained on the remaining datasets. The held-out genome was segmented into overlapping 16,384 bp windows with a 1,024 bp stride. For each window, we averaged the predictions from all models in the ensemble to produce a final predicted RNA-seq coverage profile. We quantified prediction performance using the median Spearman correlation coefficient between the predicted and observed coverage tracks over all test windows.

#### 2.7.3. Comparison to Yorzoi

To enable a fair comparison between ExoShorkie and Yorzoi, the only current method for RNA-seq coverage prediction of exogenous genomes in yeast, we modified its prediction pipeline to account for differences in input length, output resolution, and the number of predicted tracks. Yorzoi operates on input windows of 4,992 bp and predicts coverage over the central 3,000 bp at a 10 bp resolution. We first applied Yorzoi to the full test sequences using a stride of 3,000 bp, corresponding to its predicted region, to generate genome-wide predictions for all 162 output tracks.

We then unbinned these predictions and inverse-transformed them using Yorzoi’s built-in utilities to recover coverage values at single-nucleotide resolution. We subsequently averaged Yorzoi’s predicted single-nucleotide coverage across the 10 exogenous human RNA-seq output tracks to obtain a single aggregated prediction for exogenous DNA, which we refer to as *ExoYorzoi*. We segmented the resulting genome-wide predictions into overlapping windows matching ExoShorkie’s input length using a stride of 1,024 bp. For each window, we extracted the central 14,336 bp region and summed the predicted coverage into non-overlapping 16 bp bins, yielding an 896-dimensional prediction vector. Finally, we computed the median Spearman correlation coefficient between ExoYorzoi’s predictions and the corresponding experimentally measured RNA-seq coverage across all test windows.

### 2.8. Interpretability analysis

#### 2.8.1. ExoShorkie model distillation

To enable *in silico* mutagenesis (ISM) with a single model, we distilled each dataset-specific ExoShorkie ensemble into a student network trained to match the ensemble’s average prediction.

For each dataset, we first constructed a base set of approximately 10,000 overlapping input sequences using our dataset-specific stride strategy. We then generated augmented training examples using a sequence-augmentation pipeline adapted from AlphaGenome [Avsec et al., 2025]. Specifically, for each sampled sequence we applied: (i) reverse-complement augmentation with probability 0.5, (ii) random point mutations affecting 4% of positions, and (iii) structural variations consisting of random insertions, deletions, and inversions. The number of structural variation events per sequence was drawn from a Poisson distribution with *λ* = 1, and the length of each variation was sampled uniformly from [1, 20] bp.

We created 50,000 synthetic training sequences by sampling from the base set of sequences with replacement and applying the augmentation pipeline. We then obtained training targets for the student model by running the corresponding 40-model ensemble on each synthetic sequence and averaging the predicted profiles across ensemble members.

The student model used the same architecture as ExoShorkie and was initialized from the trained Shorkie fold-0 weights, with the task-specific output layer randomly initialized. We trained the student model for 20 epochs with a batch size of 64 using the Adam optimizer (default parameters, consistent with ExoShorkie training) and a learning rate of 2 *·*10^−5^. After training, the student’s predictions were highly correlated with the ensemble outputs (Pearson *r >* 0.98 on the distillation set).

#### 2.8.2. In silico mutagenesis analysis of ExoShorkie

We implemented an ISM pipeline adapted from the Shorkie model [Chao et al., 2025]. Given the lack of prior regulatory annotations for the exogenous sequences, we performed systematic mutagenesis of every nucleotide across each exogenous genome. We utilized a sliding window with a stride of 14,336 bp to ensure that each position was located within the prediction region of the model at least once.

For each window, we substituted every base with the three alternative nucleotides to generate coverage predictions by the model. We inverse-transformed these raw log Z-score outputs to their original coverage scale using the training-derived mean and standard deviation, and summed the predicted coverage values across all output bins for both the mutated and reference sequences. We then defined the ISM score matrix *M*_*i,n*_, where each entry corresponds to the log_2_ fold-change in predicted coverage obtained by substituting the reference nucleotide at genomic position *i* with nucleotide *n*:

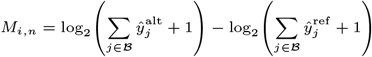

where 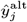 and 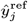 denote the model-predicted coverage for genomic bin *j* for the mutated and reference sequences, respectively, and *B* indexes the set of output bins (896 bins) produced by the model for a given window. We next centered the attribution scores by subtracting the mean over the four nucleotides at each position:

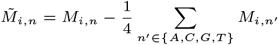

To focus exclusively on reference-base contributions, we computed:

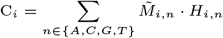

where *H*_*i,n*_ ∈ {0, 1} denotes the one-hot indicator of the reference nucleotide at position *i*. This procedure yields a vector of nucleotide-level contribution scores across the sequence, which we use for visualization and subsequent motif discovery for each exogenous genome (Fig. 2).

**Fig. 2.**
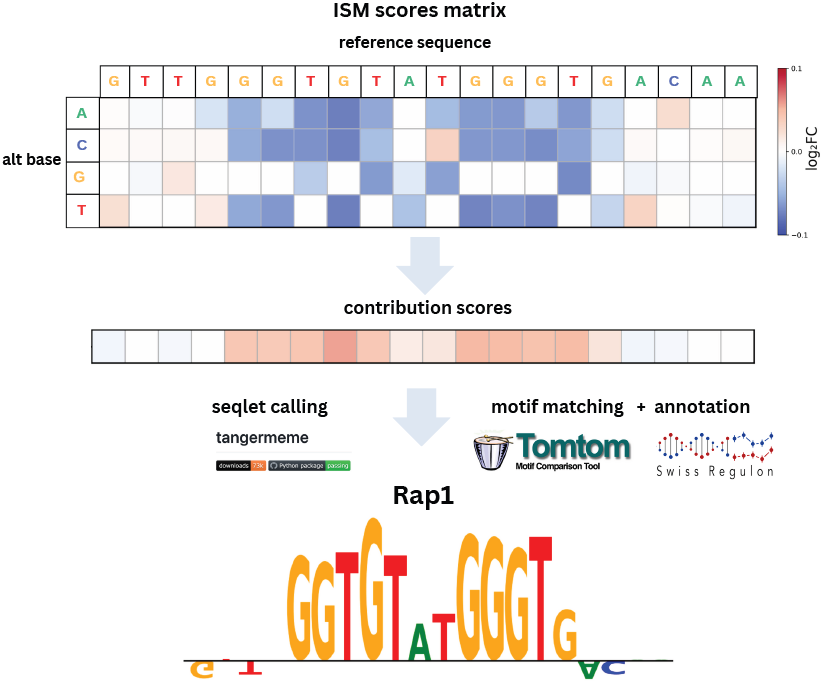
ISM pipeline schematic. For each window, each nucleotide position was substituted with the three alternative bases, and an ISM score matrix was computed as the log2 fold-change (log2FC) between the summed predicted coverage of the alternative and reference sequences. At each nucleotide position, the mean log2FC across the four nucleotides was subtracted, and reference-base contribution scores were extracted. These nucleotide-level contribution scores were subsequently used for seqlet calling and motif matching against the SwissRegulon database [Pachkov et al., 2012].

#### 2.8.3. Seqlet calling and motif identification

We utilized the nucleotide-level contribution scores generated by the ISM pipeline to identify potential regulatory regions within the attribution maps. To identify seqlets with high contribution scores, we applied the recursive seqlet calling algorithm from the tangermeme toolkit [Schreiber, 2025a]. We executed the seqlet-calling algorithm with a minimum seqlet length of 6 bp, a maximum seqlet length of 25 bp, and a p-value threshold of 0.001.

Following the identification of high-importance seqlets, we matched them to a motif database using the same tangermeme toolkit. The tangermeme toolkit leverages tomtom-lite [Schreiber, 2025b], an optimized implementation of the Tomtom motif similarity algorithm [Gupta et al., 2007], to calculate alignments between one-hot encoded seqlets and a database of known position weight matrices. For this analysis, we used the SwissRegulon database, which contains 158 curated yeast-specific transcription-factor (TF) motifs [Pachkov et al., 2012]. We retained seqlets that had significant matches to motifs below an E-value threshold of 0.05.

## 3. Results

### 3.1. Native-genome-trained models achieve low prediction performance on exogenous genomes

We first established the baseline performance of models trained exclusively on native genomic sequences when evaluated on out-of-distribution, exogenous DNA. Since we fine-tune Shorkie on exogenous data, we used the trained Shorkie model, after adapting it for strand-resolved RNA-seq coverage prediction, as our native-genome-trained baseline, which we term *NatShorkie*.

We assessed NatShorkie’s performance by evaluating it on the same 5-fold data splits used for ExoShorkie and averaging the per-fold median Spearman correlations.

As expected for this out-of-distribution task, the median Spearman correlations were relatively low (Fig. 3A): between 0.3 and 0.6 on evolutionary-constrained exogenous genomes, and even lower for the data storage chromosome (0.05), which was never shaped by evolutionary selection. This behavior is consistent with prior work showing that Enformer [Avsec et al., 2021], a sequence-to-function model trained on native genomic DNA, generalizes poorly when evaluated on rearranged or synthetic regulatory elements introduced into endogenous loci [Ribeiro-dos Santos and Maurano, 2025].

**Fig. 3.**
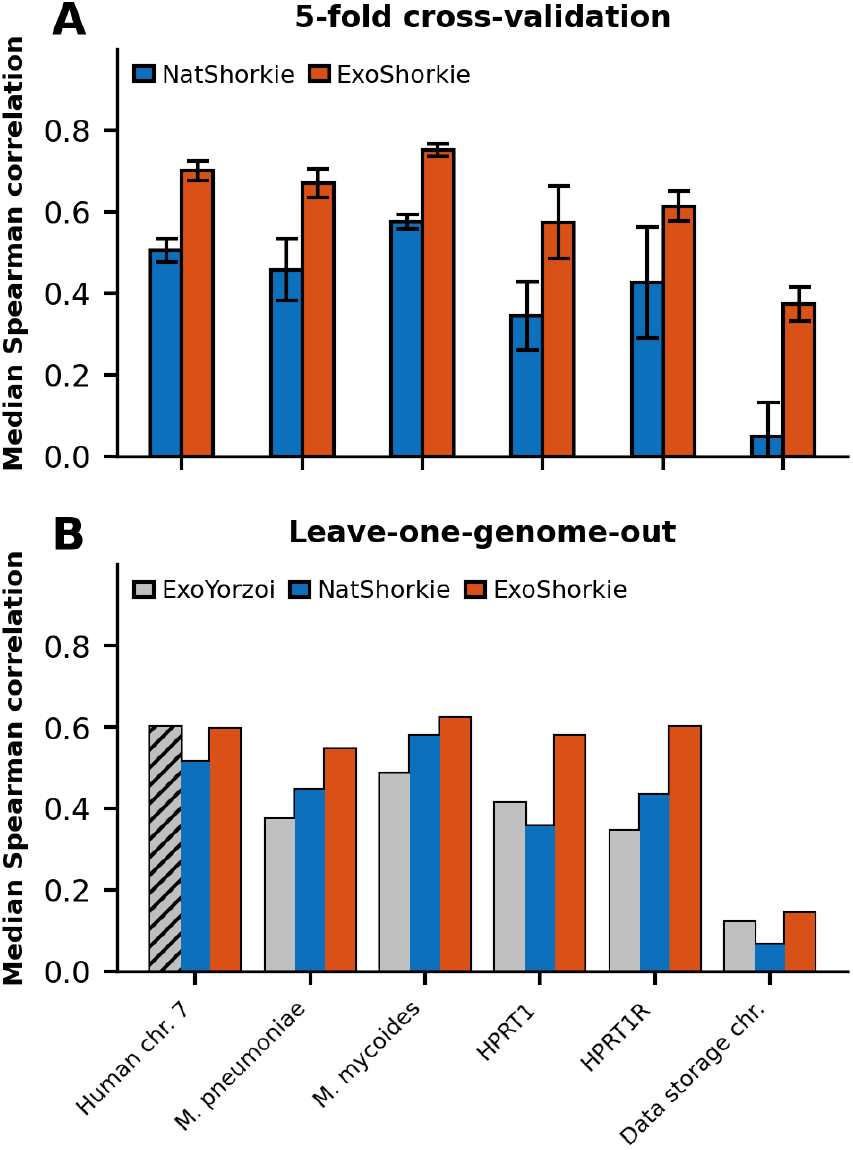
Prediction performance on exogenous genomes. (A) Comparison of median Spearman correlation over test windows between the native-genome-trained NatShorkie and ExoShorkie in 5-fold cross-validation. Error bars indicate the standard deviation across the five cross-validation test folds. (B) Comparison of median Spearman correlation over test windows between ExoYorzoi, NatShorkie, and ExoShorkie. Yorzoi was partially trained on human chromosome 7, whereas ExoShorkie was not exposed to this sequence.

### 3.2. ExoShorkie achieves high predictive performance in cross-validation

We next tested whether fine-tuning NatShorkie, the native-genome-trained baseline, on exogenous DNA sequences, which we term ExoShorkie, could improve its performance on held-out sequences from the same exogenous source. We applied a 5-fold cross-validation strategy, partitioning each exogenous sequence into 5 parts based on genomic coordinates to avoid train-test overlaps. Cross-validation significantly improved performance on all datasets compared to NatShorkie, with the average median Spearman correlation across the 5 folds increasing by 0.17–0.32 for all studies (Fig. 3A, one-sided Wilcoxon signed-rank test on per-genome averages of per-fold median Spearman correlations, *p* = 0.016). The highest increase was observed in the data storage chromosome, where the median Spearman correlation improved from 0.05 *±* 0.08 in the native baseline to 0.37 *±* 0.04. This demonstrates that ExoShorkie was able to learn regulatory rules where the native-genome-trained baseline NatShorkie failed, despite the sequence lacking evolutionary history.

### 3.3. ExoShorkie generalizes to unseen exogenous genomes

To evaluate ExoShorkie’s ability to generalize to exogenous genomes not observed during training, we performed a leave-one-genome-out evaluation. In this setting, in each iteration, one exogenous genome was held out as a test set, while predictions were generated by averaging the outputs of all ensemble models trained on the remaining genomes. This task is fundamentally more challenging than cross-validation within a single study, as the model cannot rely on genome- or experiment-specific biases and must instead capture intrinsic rules underlying the expression of exogenous DNA in yeast cells.

Across all held-out genomes, ExoShorkie consistently achieved higher prediction performance than the native-genome-trained baseline NatShorkie (Fig. 3B, one-sided Wilcoxon signed-rank tests on per-window Spearman correlations, all *p <* 10^−13^). These results indicate that ExoShorkie learns generalizable regulatory principles of exogenous gene expression that transfer to previously unseen genomes. Importantly, this evaluation setting closely reflects practical applications of ExoShorkie, where the goal is to predict the expression of newly introduced exogenous sequences for which no experimental measurements are available.

### 3.4. ExoShorkie demonstrates improved generalization compared to ExoYorzoi

We next compared ExoShorkie’s performance on unseen exogenous genomes to ExoYorzoi, the aggregated exogenous prediction derived from Yorzoi [Schneider et al., 2025]. Specifically, we evaluated ExoShorkie under a leave-one-genome-out setting and compared its performance to ExoYorzoi on held-out exogenous genomes.

Across all held-out genomes, ExoShorkie achieved higher or comparable median Spearman correlations relative to ExoYorzoi (Fig. 3B). ExoShorkie significantly outperformed ExoYorzoi on four out of six held-out genomes (one-sided Wilcoxon signed-rank tests on per-window Spearman correlations, all *p <* 10^−27^). On human chromosome 7, both models showed comparable performance, with ExoYorzoi achieving a slightly higher correlation. However, this sequence was partially observed during ExoYorzoi’s training, whereas it was completely unseen by ExoShorkie in this evaluation. On the data storage chromosome, both models showed low performance, with only marginal differences between them and slightly higher correlations for ExoShorkie. This likely reflects the fully synthetic nature of the sequence, which lacks evolutionary constraints and was not shaped by natural selection.

Importantly, aside from human chromosome 7, all comparisons were performed on genomes that were entirely unseen by both models and therefore provide a direct evaluation of generalization to novel exogenous DNA. ExoShorkie demonstrates stronger generalization to new genomes than ExoYorzoi, consistent with its training on a more diverse set of exogenous genomes, whereas ExoYorzoi was trained exclusively on exogenous human DNA expressed in yeast.

### 3.5. Pretraining outperforms random initialization

To assess the contribution of pretraining to ExoShorkie’s prediction performance, we compared ExoShorkie models initialized from Shorkie’s trained weights with models of the same architecture initialized with random weights. We evaluated both models using the same 5-fold cross-validation setting.

Across all exogenous genomes, initialization from Shorkie’s trained weights achieved higher prediction performance compared to random initialization (Fig. 4, one-sided Wilcoxon signed-rank test on per-genome averages of per-fold median Spearman correlations, *p* = 0.016). These results demonstrate that pretraining on native genomic sequences provides a substantial benefit for ExoShorkie’s downstream prediction performance, consistent with the improvements reported for Shorkie in the task of RNA-seq coverage prediction on the native genome in the original study.

**Fig. 4.**
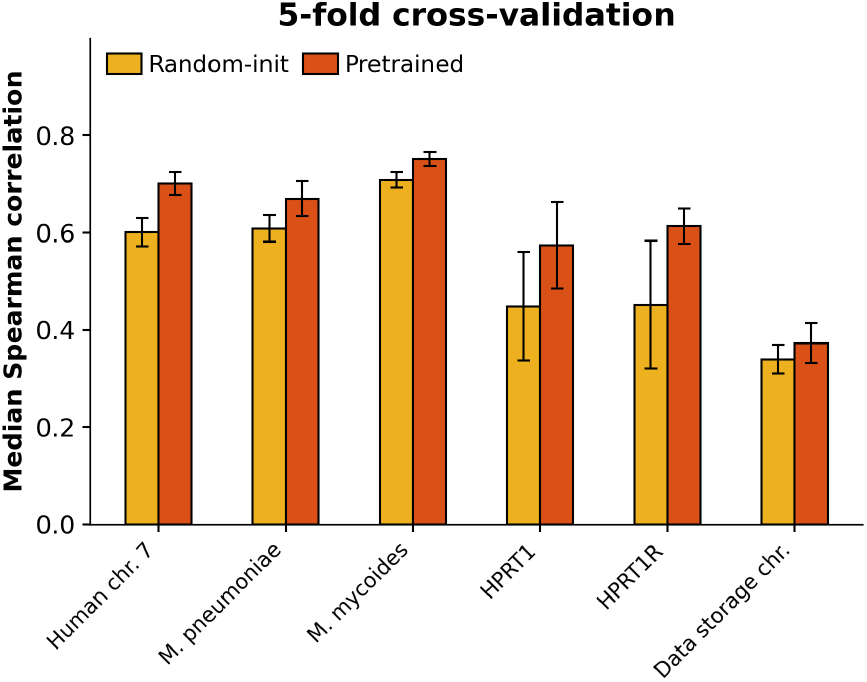
Comparison of median Spearman correlation between a pretrained ExoShorkie model, initialized with Shorkie weights, and a randomly initialized model (Random-init) with the same architecture, evaluated using the same 5-fold cross-validation setting. Performance is computed over test windows for each exogenous dataset. Error bars indicate the standard deviation across the five cross-validation test folds.

### 3.6. Attribution maps of ExoShorkie recover known yeast TF motifs in exogenous DNA

To investigate the regulatory rules learned by ExoShorkie in exogenous sequences, we performed ISM on both NatShorkie and ExoShorkie. For each exogenous genome, we computed attribution maps by performing single-nucleotide substitutions at every genomic position and measuring their effects on RNA-seq coverage predictions. We used these attribution maps to identify short genomic regions with high importance for model predictions, referred to as *seqlets*, which we subsequently matched against a yeast TF motif database. ExoShorkie recovered canonical yeast regulatory motifs, including Reb1, Fkh2, Cbf1, and the TATA-box motif, within exogenous DNA, indicating that its predictions are driven by native yeast regulatory logic acting on non-native sequences.

To focus on regions where NatShorkie and ExoShorkie disagreed in their learned regulatory logic, we computed the Pearson correlation between their contribution scores over the same windows (Fig. 5). This window-based analysis allowed us to identify local regions with low agreement, highlighting candidate regulatory elements that are interpreted differently by the two models.

**Fig. 5.**
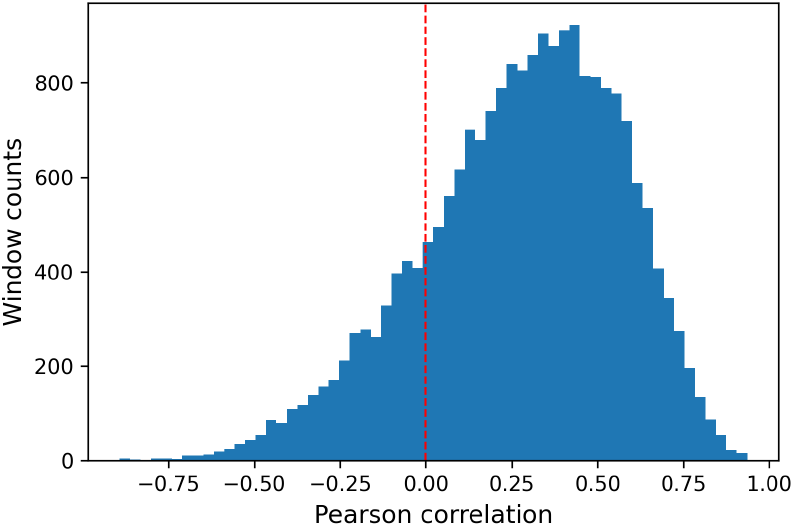
Distribution of Pearson correlations between contribution scores from ExoShorkie and NatShorkie across overlapping 150 bp windows (75 bp stride) in the *M. pneumoniae* genome.

We then examined windows in which the Pearson correlation between contribution scores from ExoShorkie and NatShorkie was negative. Within these low-agreement regions, we observed canonical yeast TF motifs that were recovered by ExoShorkie but missed by NatShorkie (Fig. 6). For windows containing these motifs, we computed the Spearman correlation between predicted and measured RNA-seq coverage and observed substantially higher correlations for ExoShorkie compared to NatShorkie. We applied the same interpretability pipeline across all exogenous genomes and observed similarly that ExoShorkie recovers exogenous-specific elements which NatShorkie missed (Supplementary Figs. S1-S5).

**Fig. 6.**
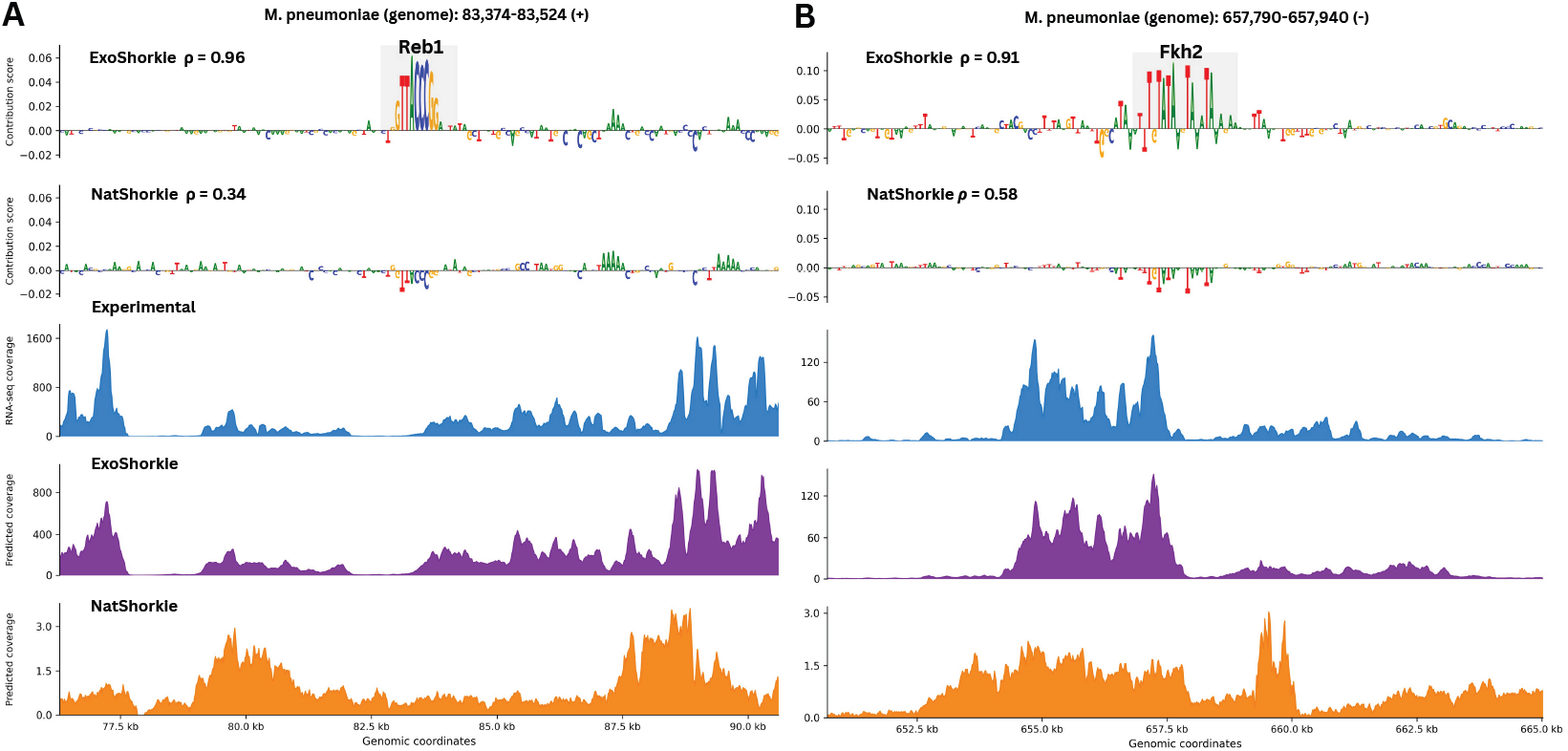
ExoShorkie recovers canonical yeast TF motifs from ISM attribution maps on the *M. pneumoniae* exogenous genome that are not recovered by NatShorkie. (A) Contribution scores from ExoShorkie (top) recover a Reb1 motif that is absent in NatShorkie (bottom). The Spearman correlation (*ρ*) between predicted and measured RNA-seq coverage within windows centered on the motif is significantly higher for the ExoShorkie model. ExoShorkie’s predicted coverage profile is more concordant with the true RNA-seq signal across the central 14,336 bp predicted region of the 16,384 bp window centered on the motif. (B) Contribution scores from ExoShorkie (top) recover a Forkhead (Fkh2) motif that is missed by NatShorkie (bottom). The Spearman correlation between predicted and measured coverage in motif-centered windows is significantly higher for ExoShorkie, and the predicted coverage profile is also more concordant with the true RNA-seq signal.

Together, these results demonstrate that ExoShorkie learns regulatory logic that differs from NatShorkie when modeling exogenous DNA and that this difference in regulatory rules translates into improved predictive performance.

## 4. Discussion

In this study, we demonstrated through ExoShorkie that combining transfer learning over the native yeast genome with fine-tuning on diverse exogenous RNA-seq datasets substantially improves performance on exogenous RNA-seq coverage prediction. By leveraging Shorkie, a pretrained state-of-the-art yeast expression model, and fine-tuning it on multiple recently published RNA-seq datasets from exogenous genomes, ExoShorkie achieves state-of-the-art performance in predicting the expression of held-out exogenous DNA sequences, outperforming both a native-genome-trained baseline and Yorzoi, which is the only existing exogenous RNA-seq coverage prediction method. Our results demonstrate that training on diverse exogenous genomes from multiple sources substantially improves generalization to new exogenous genomes. Overall, transfer learning from a yeast expression model consistently improves prediction performance across all evaluated exogenous genomes.

Our interpretability analysis via ISM shows that ExoShorkie captures canonical yeast TF motifs, including Reb1, Fkh2, Cbf1, and the TATA-box, in exogenous sequences, despite the fact that these sequences were not constrained by yeast evolution. Moreover, comparison of ISM attribution maps in regions where ExoShorkie and the native-genome-trained baseline NatShorkie disagree revealed distinct regulatory motifs identified by ExoShorkie and missed by NatShorkie (Fig. 6). These regions correspond to windows in which ExoShorkie achieves substantially higher concordance with the ground-truth RNA-seq profile compared to NatShorkie, suggesting that the distinct regulatory logic learned by ExoShorkie is the reason for its improved prediction performance.

There are several limitations to our study. First, our analysis is constrained by the small number of studies that have published RNA-seq data of exogenous genomes in yeast. Despite recent advances in synthetic genomics, the synthesis and delivery of exogenous DNA remain labor- and resource-intensive, which restricts the availability of large and diverse datasets. We expect that future technological developments will help expand the scale of exogenous RNA-seq data that can be used to train ExoShorkie. Second, our study is currently limited to a yeast host system. While yeast is a widely used model organism for studying gene regulation, the regulatory rules learned by ExoShorkie may be yeast-specific and may not directly generalize to other host organisms. Third, ExoShorkie achieves its lowest performance on highly artificial data storage DNA. Sequences that lack known evolutionary or designed regulatory elements may represent a more challenging prediction task and require substantially larger and more diverse training datasets than are currently available.

There are several directions in which we plan to extend ExoShorkie in the future. First, experimental validation of ExoShorkie’s model-identified regulatory elements would strengthen the evidence for ExoShorkie’s ability to recover exogenous-specific elements. Second, we plan to extend the model to jointly predict multiple genomic and epigenomic tracks by incorporating datasets from additional assays, such as ATAC-seq and ChIP-seq, measured on native and/or exogenous genomes. Finally, we plan to generalize our approach to exogenous RNA-seq experiments in mammalian systems [Brosh et al., 2023], in combination with recent mammalian sequence-to-expression models [Avsec et al., 2021, Linder et al., 2025].

## 5. Conclusion

In conclusion, we introduced ExoShorkie, an interpretable method for predicting RNA-seq coverage from exogenous DNA sequences that generalizes across unseen genomes and outperforms existing methods. Our results show that ExoShorkie learns regulatory patterns that differ between native and exogenous sequence contexts. Beyond prediction, ExoShorkie provides a foundation for the rational design of long multi-gene synthetic DNA constructs with predictable expression levels by sequence optimization methods, such as Ledidi [Schreiber et al., 2025]. In addition, ExoShorkie offers a general framework for studying the transcriptional regulation of exogenous sequences in yeast, advancing our understanding of gene regulation beyond native genomic contexts.

## Acknowledgments

J.M. acknowledges the Bar-Ilan University Presidential Doctoral Fellowship and the cloud computing credit by the Israel Data Science and AI Initiative.

## Data availability

The reference exogenous genome sequences and processed RNA-seq datasets used in this study are available on Figshare at https://doi.org/10.6084/m9.figshare.31075375. Trained ExoShorkie models are available on Hugging Face at https://huggingface.co/Jonathan-Mandl/ExoShorkie-models.

