## Supplementary Information for "ExoShorkie: Predicting RNA-seq coverage of exogenous genomes in yeast by transfer learning"

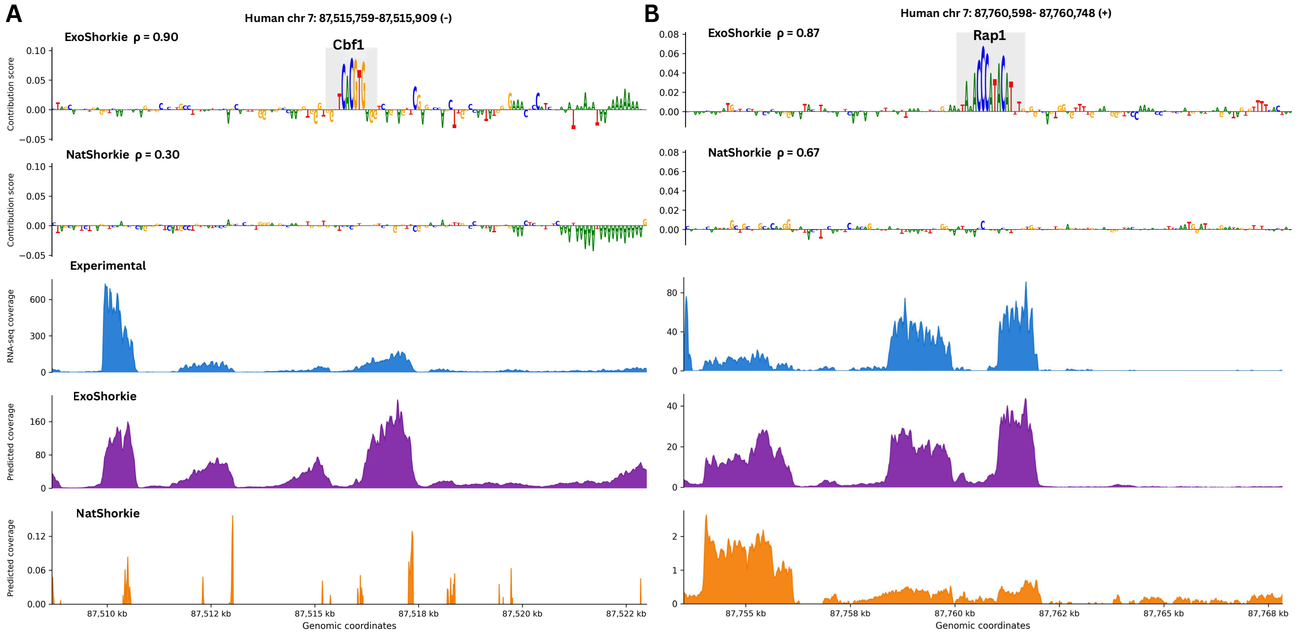


**Fig. S1.** ExoShorkie recovers canonical yeast TF motifs from ISM attribution maps on the exogenous human chromosome 7 that are diffuse or absent in NatShorkie. (A) Contribution scores from ExoShorkie (top) recover a Cbf1 motif that is not recovered by NatShorkie (bottom). (B) Contribution scores from ExoShorkie (top) recover a Rap1 motif that is missed by NatShorkie (bottom). For both loci, the Spearman correlation ($\rho$) between predicted and measured RNA-seq coverage in motif-centered windows is significantly higher for ExoShorkie. ExoShorkie's predicted coverage profile is more concordant with the true RNA-seq signal across the central 14,336 bp predicted region of the 16,384 bp window centered on each motif.

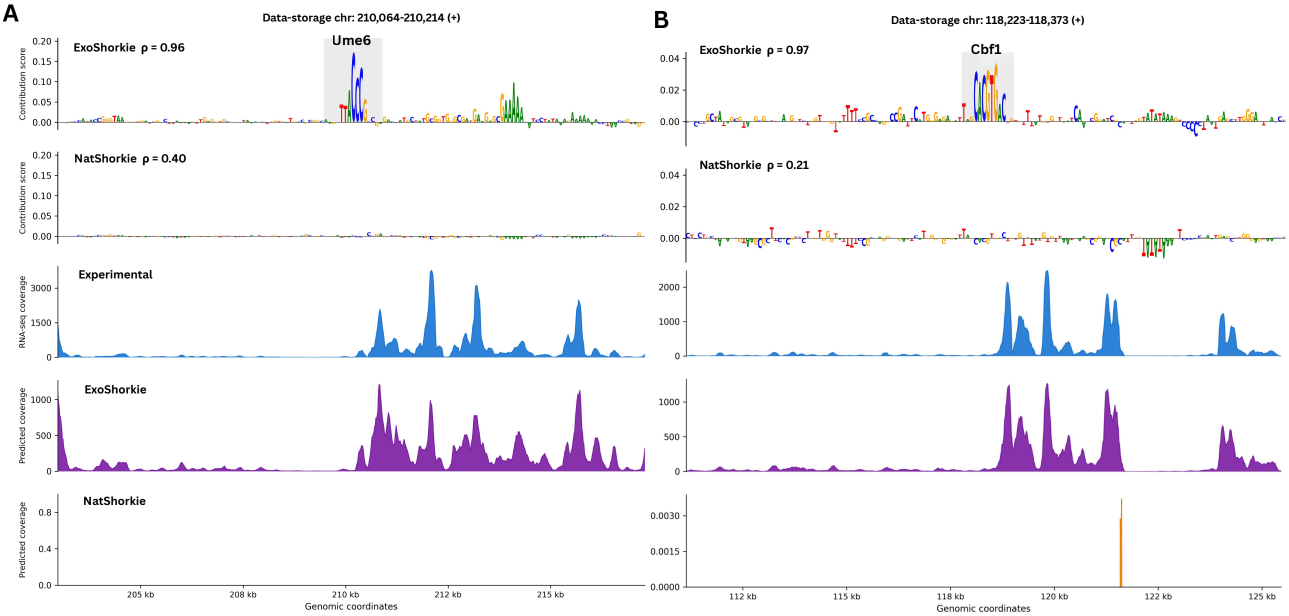


**Fig. S2.** ExoShorkie recovers canonical yeast TF motifs from ISM attribution maps on the exogenous data storage chromosome that are diffuse or absent in NatShorkie. (A) Contribution scores from ExoShorkie (top) recover a Ume6 motif that is not recovered by NatShorkie (bottom). (B) Contribution scores from ExoShorkie (top) recover a Cbf1 motif that is missed by NatShorkie (bottom). For both loci, the Spearman correlation ($\rho$) between predicted and measured RNA-seq coverage in motif-centered windows is significantly higher for ExoShorkie. ExoShorkie's predicted coverage profile is more concordant with the true RNA-seq signal across the central 14,336 bp predicted region of the 16,384 bp window centered on each motif.

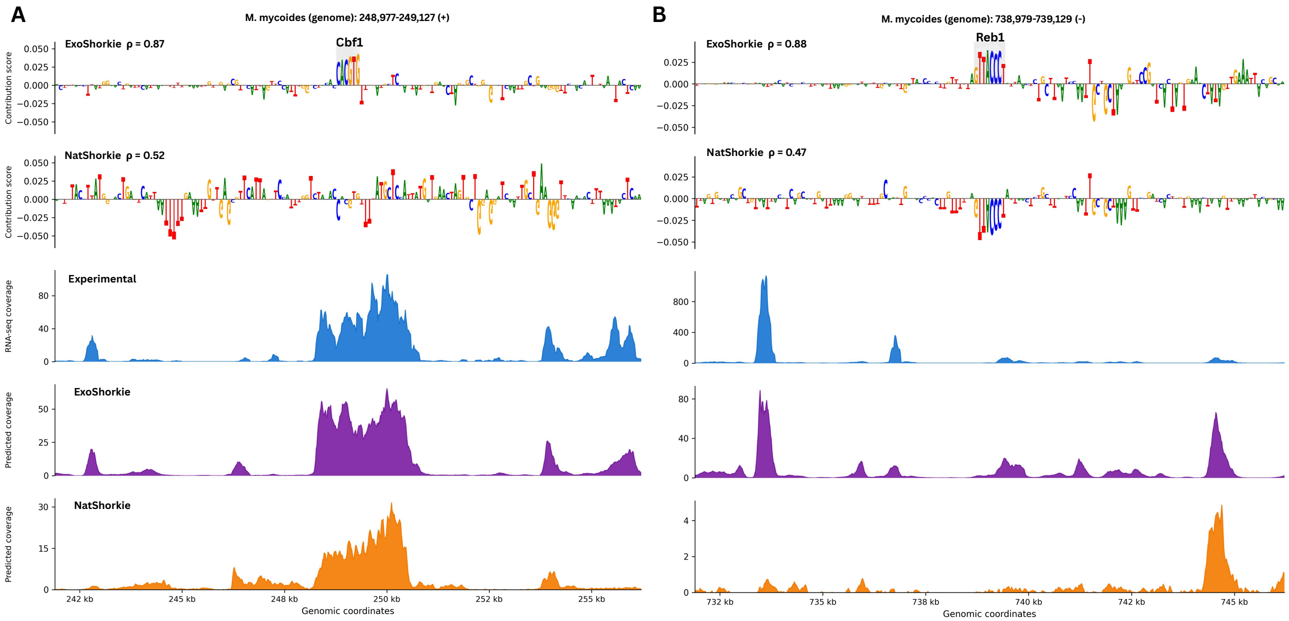


**Fig. S3.** ExoShorkie recovers canonical yeast TF motifs from ISM attribution maps on the exogenous *M. mycoides* genome that are diffuse or absent in NatShorkie. (A) Contribution scores from ExoShorkie (top) recover a Cbf1 motif that is not recovered by NatShorkie (bottom). (B) Contribution scores from ExoShorkie (top) recover a Reb1 motif that is not recovered by NatShorkie (bottom). For both loci, the Spearman correlation ($\rho$) between predicted and measured RNA-seq coverage in motif-centered windows is significantly higher for ExoShorkie. ExoShorkie's predicted coverage profile is more concordant with the true RNA-seq signal across the central 14,336 bp predicted region of the 16,384 bp window centered on each motif.

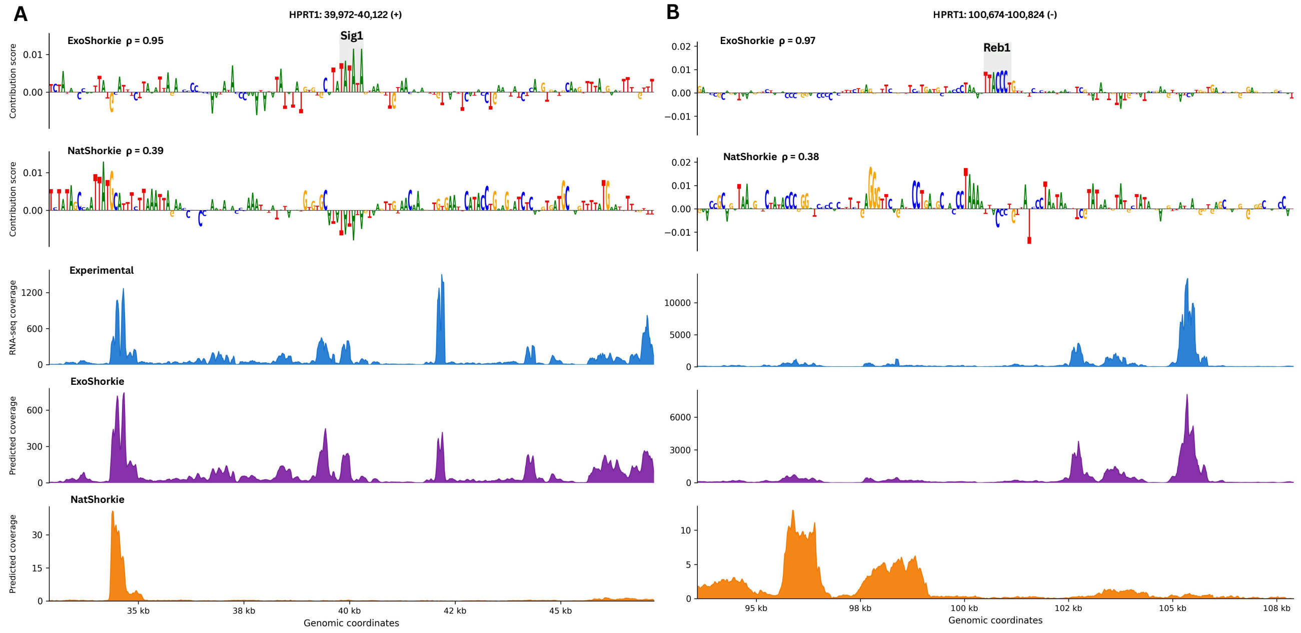


**Fig. S4.** ExoShorkie recovers yeast TF motifs from ISM attribution maps on the exogenous HPRT1 genome that are diffuse or absent in NatShorkie. (A) Contribution scores from ExoShorkie (top) recover a Sig1 motif that is not recovered by NatShorkie (bottom). (B) Contribution scores from ExoShorkie (top) recover a Reb1 motif that is not recovered by NatShorkie (bottom). For both loci, the Spearman correlation ($\rho$) between predicted and measured RNA-seq coverage in motif-centered windows is significantly higher for ExoShorkie. ExoShorkie's predicted coverage profile is more concordant with the true RNA-seq signal across the central 14,336 bp predicted region of the 16,384 bp window centered on each motif.

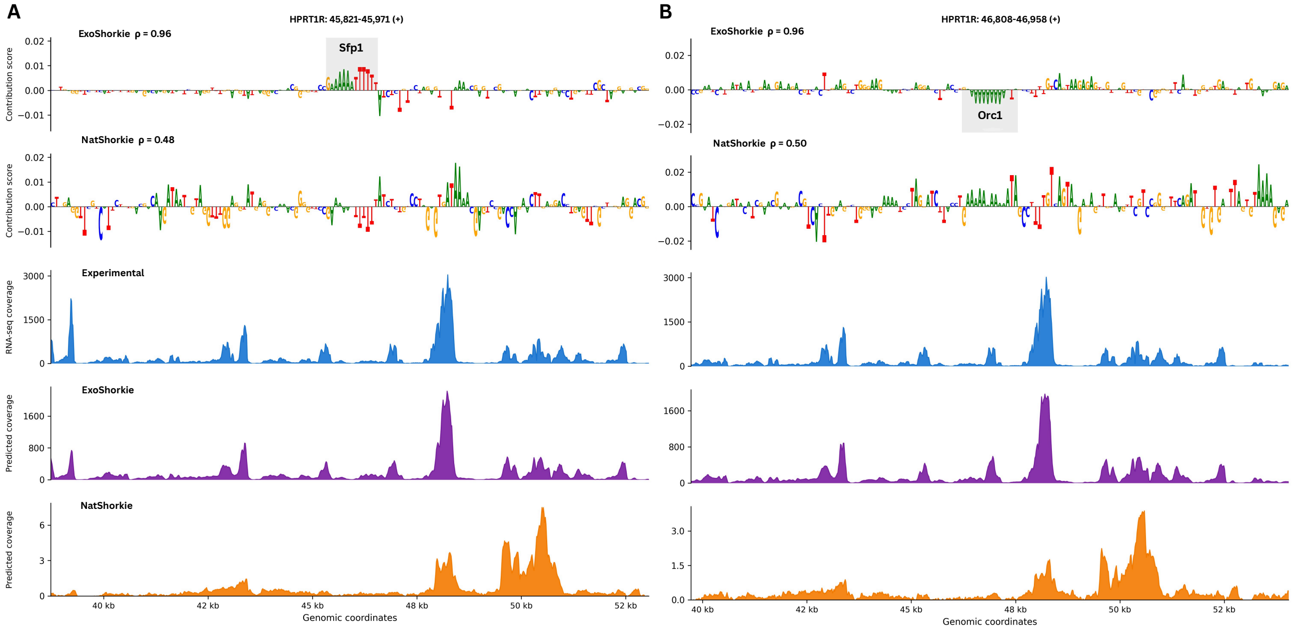


**Fig. S5.** ExoShorkie recovers yeast TF motifs from ISM attribution maps on the exogenous HPRT1R genome that are diffuse or absent in NatShorkie. (A) Contribution scores from ExoShorkie (top) recover a Sfp1 motif that is not recovered by NatShorkie (bottom). (B) Contribution scores from ExoShorkie (top) recover a Orc1 motif that is not recovered by NatShorkie (bottom). For both loci, the Spearman correlation ($\rho$) between predicted and measured RNA-seq coverage in motif-centered windows is significantly higher for ExoShorkie. ExoShorkie's predicted coverage profile is more concordant with the true RNA-seq signal across the central 14,336 bp predicted region of the 16,384 bp window centered on each motif.
